# Hidden genomic diversity of SARS-CoV-2: implications for qRT-PCR diagnostics and transmission

**DOI:** 10.1101/2020.07.02.184481

**Authors:** Nicolae Sapoval, Medhat Mahmoud, Michael D. Jochum, Yunxi Liu, R. A. Leo Elworth, Qi Wang, Dreycey Albin, Huw Ogilvie, Michael D. Lee, Sonia Villapol, Kyle M. Hernandez, Irina Maljkovic Berry, Jonathan Foox, Afshin Beheshti, Krista Ternus, Kjersti M. Aagaard, David Posada, Christopher E. Mason, Fritz Sedlazeck, Todd J. Treangen

## Abstract

The COVID-19 pandemic has sparked an urgent need to uncover the underlying biology of this devastating disease. Though RNA viruses mutate more rapidly than DNA viruses, there are a relatively small number of single nucleotide polymorphisms (SNPs) that differentiate the main SARS-CoV-2 clades that have spread throughout the world. In this study, we investigated over 7,000 SARS-CoV-2 datasets to unveil both intrahost and interhost diversity. Our intrahost and interhost diversity analyses yielded three major observations. First, the mutational profile of SARS-CoV-2 highlights iSNV and SNP similarity, albeit with high variability in C>T changes. Second, iSNV and SNP patterns in SARS-CoV-2 are more similar to MERS-CoV than SARS-CoV-1. Third, a significant fraction of small indels fuel the genetic diversity of SARS-CoV-2. Altogether, our findings provide insight into SARS-CoV-2 genomic diversity, inform the design of detection tests, and highlight the potential of iSNVs for tracking the transmission of SARS-CoV-2.

## Introduction

Coronavirus (CoV) genomes are the largest among single strand RNA (ssRNA) viruses, ranging from 26 to 32 Kbp. While ssRNA viruses typically display very high mutation rates, coronaviruses encode an RNA polymerase with 3’-to-5’ proofreading activity that allows them to replicate their genome with high-fidelity, lowering their mutation rate (*1*–*4*). Additionally, SARS-CoV-2 contains a common 69-bp 5’ leader sequence fused to the body sequence from the 3’ end of the genome (*5*). Then, leader-to-body fusion occurs during negative-strand synthesis at short motifs called transcription-regulating sequences (TRS), which are conserved 7 bp sequences that are adjacent to the ORFs.

On March 11, 2020, the WHO determined that an outbreak of a novel coronavirus SARS-CoV-2 that began in Wuhan, China in December 2019 had reached pandemic status. Initial consensus-level genomic data from the Global Initiative on Sharing All Influenza Data (GISAID) (*6*) indicated that the SARS-CoV-2 mutational rate (*7*) was similar to other CoVs (*8*). In order to properly assess the genomic diversity of any RNA virus, and specifically SARS-CoV-2, it is necessary to also consider the intrahost polymorphisms (*9*–*12*), including often overlooked structural variation. Recent studies have claimed that host-dependent RNA editing might be a key factor for understanding the mutational landscape of SARS-CoV-2 within hosts (*13, 14*). However, these studies were based on a limited number of samples (<20). In order to explore both the intrahost and interhost mutational landscape of SARS-CoV-2, we leveraged a dataset consisting of 6,928 consensus genomes from GISAID, 11 sequencing samples from the Baylor College of Medicine, and 140 sequencing samples from the Weill Cornell College of Medicine.

Understanding the intrahost genomic diversity of SARS-CoV-2 is also important for different applications. Most SARS-CoV-2 detection tests rely on oligonucleotide probes and primers that must be sensitive to SARS-CoV-2. In this setting, sensitivity determines how well it can capture the diversity of all SARS-CoV-2 variants. Lack of sensitivity leads to an increase in false positive qRT-PCR results, as few as two mismatches can result in increases in CT values and degradation in accuracy of viral load estimates (*15, 16*). Moreover, recent studies on Ebolavirus and flu viruses (*12, 17*) highlight the importance of intrahost variation for studying viral population dynamics and transmission scenarios. In summary, in this study, we investigate the intrahost diversity of SARS-CoV-2 by conducting a broad evaluation of (i) intrahost single nucleotide variants (iSNV), (ii) consensus-level single nucleotide polymorphisms (SNPs), and (iii) structural variants, across assembled genomes, amplicon, and metatranscriptomic datasets totaling over 7,000 samples.

## Results

We analyzed three SARS-CoV-2 genomic datasets: GISAID public consensus sequences, sequencing reads for 11 samples collected by the Baylor College of Medicine in Houston, and sequencing reads for 140 samples collected by Weill Cornell University in New York City (NYC). We evaluated structural variants across the 151 samples in both NYC and Houston; the inferred SVs are shown in Figure 1A. We also evaluated single nucleotide variants in GISAID representing single nucleotide polymorphisms (SNPs), while the variants analyzed in the Houston and NYC datasets include both SNPs and intrahost single nucleotide variants (iSNVs). The inferred phylogenetic tree of GISAID genomes with clade-defining (*18*) SNPs is shown in Figure 1B. We note that these previously reported clade-defining SNPs distinguish the geographic origin of SARS-CoV-2 genomes, with clades G and S predominantly covering North American genomes and clade V covering a portion of Asian and European genomes. We also observe that some of the clade-defining SNPs occur intermittently outside of the main phylogenetic clades. We will now dive deep into three main results: (i) intrahost structural variant (SV) landscape, (ii) intrahost single nucleotide variant (iSNV) landscape, and (iii) exploratory analyses of shared SNPs and iSNVs within and across patients in NYC.

**Figure 1:**
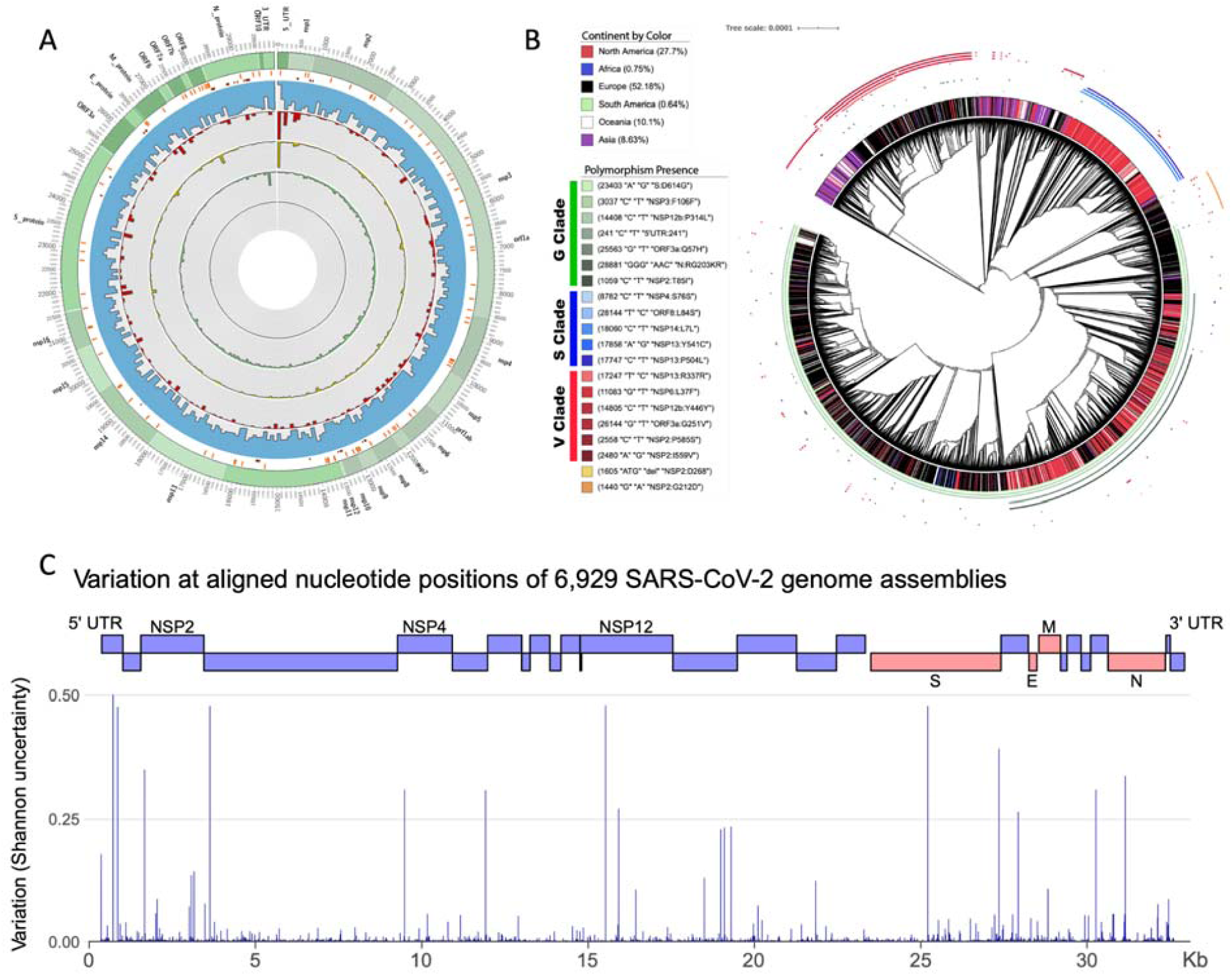
Overview of general diversity of SARS-CoV-2. **A.** From outer to inner layers: Annotation of SARS-CoV-2 genome (green), transcription-regulating sequences (TRS) (orange), PCR primer designs (dark red), intrahost variant density including iSNVs (blue), deletions start sites (red), duplication start sites (yellow), inversion start sites (green) and insertions (dark green) along the entire genome. For SNPs + iSNVs + SVs we plotted the density scaled by their allele frequency across the population over 100bp windows. **B.** Directly outside of the tree branches is the continuous annotation ring for the continents corresponding to each GISAID sample. The set of smaller non-continuous rings, surrounding the continent annotation ring, are the clade-specific SNPs as described in (*18*). The G clade SNPs are colored as different shades of green, the S clade ones are colored different shades of blue, and the V clade ones are different shades of red. **C.** This figure shows the variability of positions in SARS-CoV-2 overlaid with the protein coding regions in the genome.

### Intrahost Structural Variant (SV) Landscape

We identified 3,311 structural variants (SVs) across 170 sequencing samples, with the majority being inversions (1,504) and tandem duplications (1,157), followed by deletions (625) and a few insertions (25) (Figure 1A). Overall, since we are identifying SVs based on RNA-Seq data, the majority of these SVs are likely to be highlighting variability in the SARS-CoV-2 transcriptome (*16*), which is influenced by fusion, deletions, frame-shifts, and recombination. We observed a significant overlap (Kolmogorov–Smirnov test: p-value=4.95^−5^, D=0.25) for the 98 start and 63 end breakpoints with the annotated transcription regulating sequences (TRS) (dark red Figure 1A). Subsequently, we focus on smaller SVs (*<*1kbp) that more likely indicate true underlying SV rather than transcription signals. We identified 247 deletions and 23 insertions across all 170 SARS-CoV-2 genomes. The imbalance of insertions and deletions is likely due to the low ability to detect insertions using short reads (*19*). Figure 1A shows the allele frequency (AF) of these SVs across all samples. We observed 8 deletions shared among 34 or more samples (AF: >20%): a 14bp at 509bp (NSP1) (AF: 30.59%), a 9bp at 685bp (NSP1) (AF: 23.53%), a 24bp at 4532 (NSP3) (AF:25.29%) a 39bp at 21740bp (spike protein) (AF: 37.65%), a 22bp at 23558bp (spike protein) (AF: 31.76%), a 15bp at 24014bp (spike protein) (AF: 21.18%), a 41bp at 26779bp (M protein) (AF: 34.12%) and a 14bp at 29067 (N protein) (AF: 20%).

Next, we investigated where these SVs are mainly located with respect to the annotated regions. We identified an enrichment of SVs in NSP11 and NSP12 when taking the size of the annotated regions into account (Supplementary Figure 1). In addition, it is interesting to see that a higher number of SVs are also clustering in E protein (5 del), NSP7 (5 del and 1 ins), NSP9 (7 del and 1 ins), ORF6 (6 del) and ORF7b (3 del).

We further compared our SV call set with previously reported single deletions reported by various groups. Davidson et al (*20*) reported a 24bp deletion in the subgenomic mRNA encoding the spike (S) glycoprotein that played a role in removing a proposed furin cleavage site from the S glycoprotein. We were able to identify this deletion (position: 25234bp), but only in 3 of our samples. However, in total we discovered six deletions shared among samples within the Spike protein. Three of them showed above with AF> 20% and the remaining at: 21984bp (9bp, AF:19.41%), 22824bp (78bp, AF: 11.76%) and at 24125bp (15bp, AF: 8.24%). We further identified five deletions, one (at 28245bp) was present in 10 samples (AF: 6%) in ORF8, a potentially important gene for viral adaptation to humans (*21*).

### Intrahost Single Nucleotide Variant (iSNV) Landscape

We considered intrahost single nucleotide variants (iSNVs) to be those with an AF between 2% and 50% in a sample. Above 50%, all single nucleotide variants were considered to be consensus-level single nucleotide polymorphisms (SNPs) as it is a common threshold for consensus-calling in genome assembly (*22, 23*). Figure 2A shows the iSNV AF distribution, with the peak occurring in the 2% to 5% range of the distribution. The predominant iSNVs observed are T>C and C>T (Figure 2B). We also note that A>G, G>A, and G>T iSNVs are common. When the distribution of iSNVs is mapped onto the SARS-CoV-2 genome, we observe that C>T is the dominant SNP in 10 out of 16 genes (Figure 2D). NSP6 and NSP10 stand out as having larger fractions of T>C iSNVs, and NSP7 has a large fraction of A>C iSNVs (Figure 2D). Additionally NSP6 and ORF3a have a high fraction of G>T SNPs, and ORF8 and M genes have a high fraction of T>C SNPs. We also identified several interesting patterns of SNP and iSNV mutational patterns within the ORFs of SARS-CoV-2. Of note, SARS-CoV-2 encodes three tandem macrodomains within non-structural protein 3 (NSP3). NPS3 is essential for SARS-CoV-2 replication and represents a promising target for the development of antiviral drugs (*24*). The NSP3 protein is also one of the most diverged regions of SARS-CoV-2 compared to SARS-CoV-1 and MERS-CoV.

**Figure 2:**
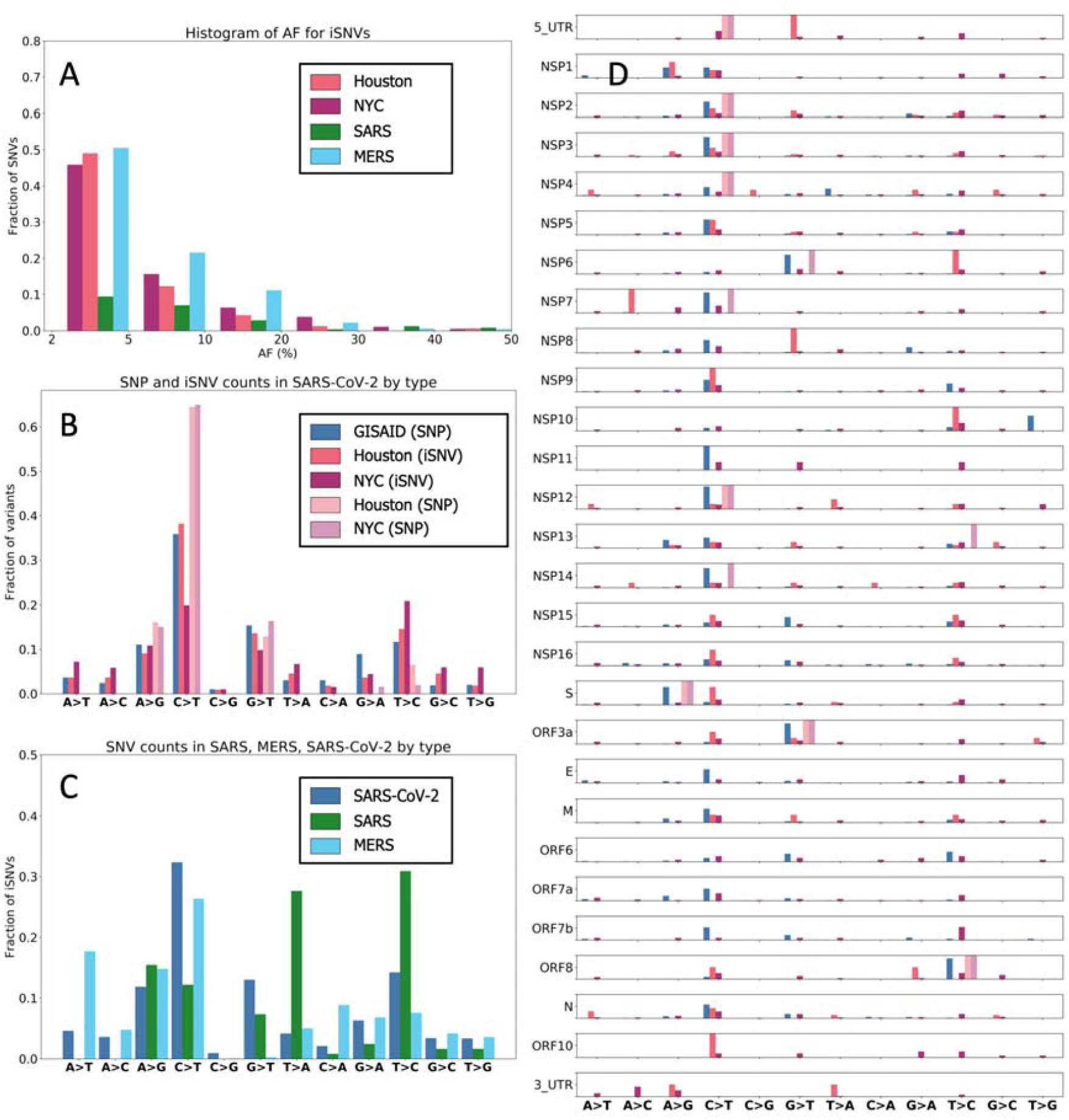
Mutational frequencies of iSNV and SNPs. **A.** *Distribution of iSNV AF.* We note that the distribution of AF is strictly less than 50% as iSNVs are below consensus-level by definition. **B.** *Mutational spectrum of SARS-CoV-2.* **C.** *Mutational spectra of SARS-CoV-1, SARS-CoV-2, and MERS.* **D.** *Mutational spectrum of SARS-CoV-2 by ORF/NSP.*

We note that the mutational spectra for SNPs matches the one observed for iSNVs, namely A>G, G>A, T>C and G>T are most common (Figure 2B). However, one striking difference is the relatively lower percentage of C>T changes in iSNVs from the NYC dataset (20%) compared to 40% C>T iSNVs for Houston samples and over 50% C>T in Houston and NYC SNPs. The fraction of GISAID C>T SNPs is nearly to the fraction of Houston C>T iSNVs, clearly distinguishing GISAID SNPs and Houston iSNVs from Houston and NYC SNPs. We also note that the mutational spectra of SNPs across the genes of SARS-CoV-2 closely match the iSNV mutational spectra (Figure 2D). The mutational spectrum of NYC SNPs is significantly different from both NYC iSNVs mutational spectrum (Kolmogorov-Smirnov (KS) test: p-value ∼ 10^−100^) and GISAID SNPs mutational spectrum (KS test: p-value ∼ 10^−40^). When compared to SARS and MERS, SARS-CoV-2 has a larger proportion of G>T iSNVs (Figure 2C). The other four major iSNV types (C>T, T>C, A>G, and G>A) are well represented in all three viruses. We also note that SARS data does not have any A>T nor A>C iSNVs.

To further investigate patterns of difference and similarity between SNPs and iSNVs, we analyzed the functional impact of the observed variants. First, in GISAID SNPs we observe 1191 (36.45%) synonymous, 2021 (61.86%) missense, and 40 (1.22%) stop gained variants. In NYC iSNVs we observed 782 (31.68%) synonymous, 1549 (62.76%) missense, and 73 (2.96%) stop gained variants. Finally, in Houston iSNVs we observed 43 (31.16%) synonymous, 86 (62.31%) missense, and 5 (3.62%) stop gained variants. Altogether, about two thirds of all observed variants are missense and about a third are synonymous, with good agreement of these values for both SNPs and iSNVs. We also investigated the overlap between iSNV and consensus-level SNPs (Figure 3B). We note that there are 15 mutations that have been found in GISAID data, NYC data, and Houston data independently. We also observed that 230 SNVs occur both as an iSNV in at least one sample and as SNPs in the GISAID data. Finally, there are 2 iSNVs that also occur as SNPs (Figure 3B). The mutational spectrum of variants that occur as both SNPs and iSNVs is similar to the general one outlined above with ∼65% of the changes being C>T, followed by ∼15% of G>T, and ∼12% of T>C.

**Figure 3:**
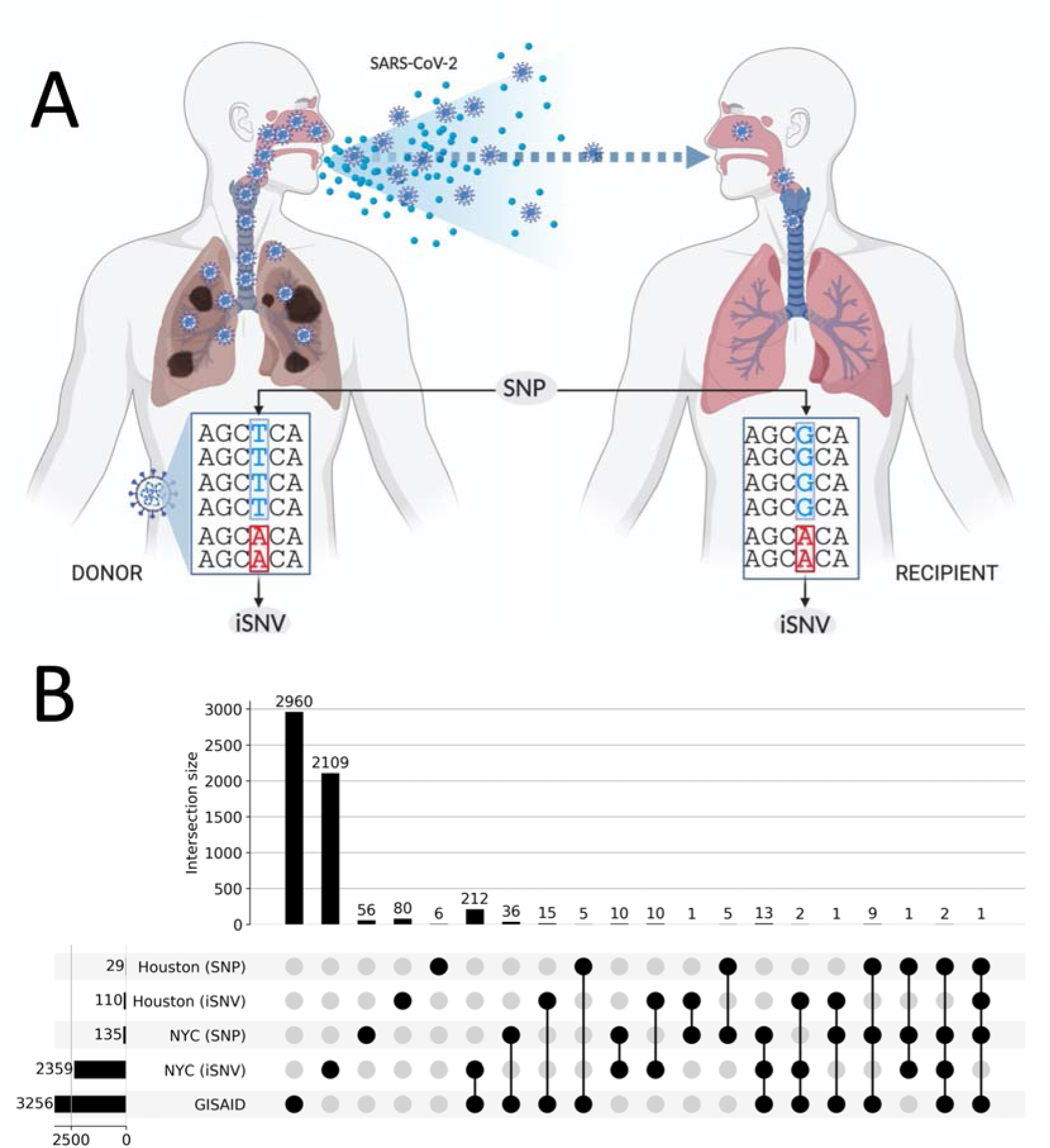
Shared SNPs and SNVs across datasets. **A.** Illustration differentiating what we define as an intrahost SNV (iSNV) and an interhost consensus-level SNP. **B**. This UpSet plot captures the shared single nucleotide variants between iSNVs and consensus-level SNPs. The horizontal bars on the left show the total number of variants in the given category. Vertical bars indicate the size of the intersection between highlighted (with black circles) sets. Every variant contributes to exactly one intersection size to avoid double counting.

Prior studies have found iSNVs early in virus outbreaks that later establish as SNPs (*25, 26*). Thus, we looked into whether clade-defining SNPs identified in a previous study (*18*) co-occur with iSNVs identified in NYC and Houston datasets. We found that a G and S clade-defining SNPs co-occur with an iSNV position 13542 in the NSP12 gene. There are two synonymous iSNVs at this position, the more common one is a T>G change (seen in both NYC and Houston), and a less common one is a T>A change occurring only in the NYC data. This indicates the emergence of an iSNV strongly correlated with the North American clade of the SARS-CoV-2.

Next, we estimated the genetic complexity (*S*_*n*_) (*27*) and genetic diversity (*π*) of SARS-CoV-2, SARS-CoV-1 and MERS (Figure 4A,B). For both diversity and complexity all three viruses show distinct distributions of (KS test: p-value < 10^−8^) with a higher variance in SARS-CoV-2. We also compared the ratios of non-synonymous and synonymous diversities (*π*_*N*_/*π*_*S*_) for iSNVs in SARS-CoV-2, SARS-CoV-1 and MERS data (Figure 4C). The genome-wide *π*_*N*_/*π*_*S*_ values suggest that SARS-CoV-2 (median *π*_*N*_/*π*_*S*_: 0.554) and SARS-CoV-1 (median *π*_*N*_/*π*_*S*_: 0.179) might be predominantly under purifying selection, while MERS (median *π*_*N*_/*π*_*S*_: 1.270) seems to be overall under positive selection (KS test: p-value < 10^−7^). We also observed a significant difference in the distribution of *π*_*N*_/*π*_*S*_ ratios between iSNVs and SNPs in the NYC data (KS test, p-value 6.29 × 10^−12^). The SARS-CoV-2 *π*_*N*_/*π*_*S*_ values are significantly lower for iSNVs (median *π*_*N*_/*π*_*S*_: 0.273) than for SNPs (median *π*_*N*_/*π*_*S*_: 0.446, Figure 4D). The *π*_*N*_/*π*_*S*_ ratios are consistent across ORFs/NSPs of SARS-CoV-2 (Supplementary Figure 2).

**Figure 4:**
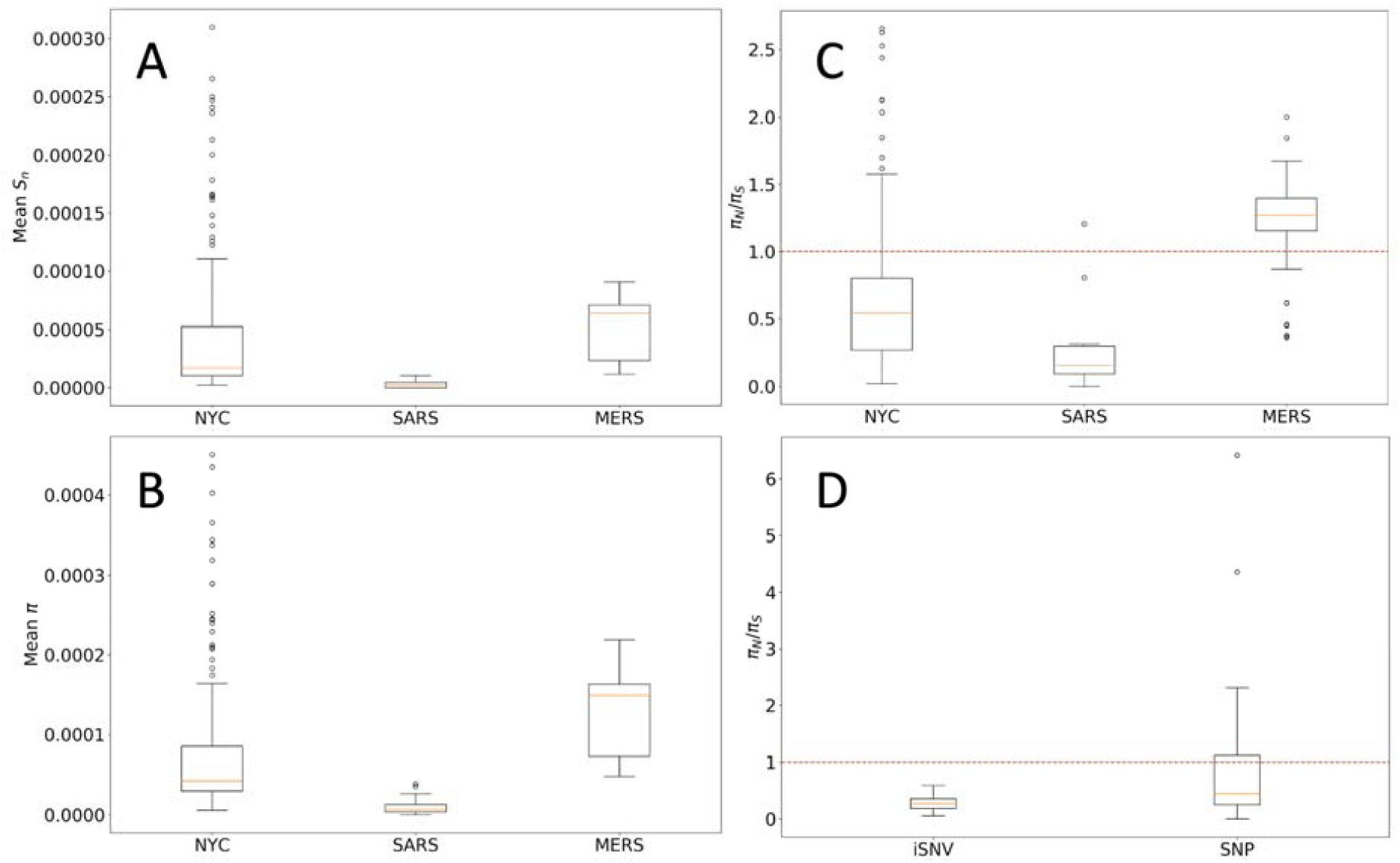
Complexity and diversity in Coronaviruses. **A.** Intrahost complexity of Coronavirus samples. This plot shows the mean *S*_*n*_ complexity of samples for SARS-CoV-2, SARS-CoV-1 and MERS. **B.** Diversity of Coronavirus samples. This plot shows the mean diversity of samples. **C.** Synonymous vs non-synonymous diversity ratios. **D.** *Syn. vs non-syn. diversity ratios for iSNVs and SNPs in NYC data.*

Finally, we analyzed the potential impact of iSNVs and SNPs on the probes and primers used for detection of SARS-CoV-2 (*15, 28*). To evaluate this, we downloaded the set of probes and primers sequences available at the WHO website, as well as the Arctic primers. Among these, 263 out of 272 sequences contained at least one SNP or iSNV (Figure 5, Table S2). On average, each probe/primer sequence contained 2.529 iSNV and/or 2.477 SNPs. These results suggest the potential for a drop in the sensitivity of the affected probes and primers. We also note that since the iSNV and SNP mutational profiles mimic each other for specific mutations, the potential impact of iSNVs on primer and probe binding should not be overlooked given the possibility of iSNVs establishing as SNPs (*26*).

**Figure 5:**
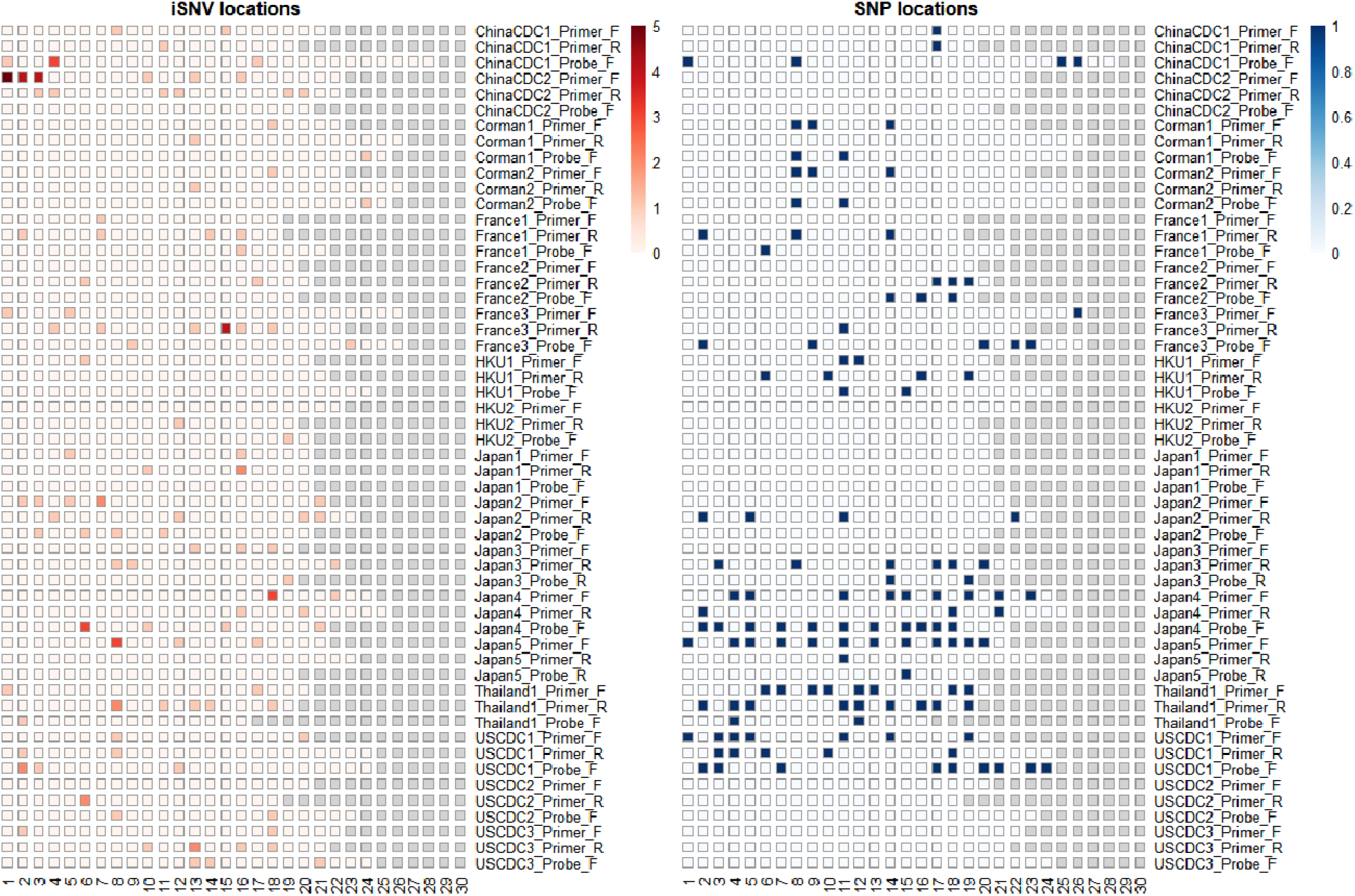
iSNV and SNP presence on widely-used primers and probes. This figure shows the locations on WHO probes and primers that contain iSNVs (left) and SNPs (right). Columns correspond to base pair positions within the probe, and the sequences are 3’ aligned. Rows corresponding to the oligonucleotide sequences and highlighted squares indicate that the position is affected by a SNV in one or more samples.

### Exploratory Transmission Analysis of Shared SNPs and iSNVs within and across patients

Shared viral genomic variants can be indicative of transmission events and routes (*29*), and iSNVs are a critically important tool for discerning direct transmission and for bottleneck calculations (*30*). To assess our ability to identify shared iSNVs and SNPs across samples, we first compared all NYC paired samples from the same patient taken within 24 hours (Figure 6A,B). In Figure 6A, we see eight shared SNPs, one shared iSNV, and two shared iSNVs that occur as a SNP in patient 340 sample C03 and as iSNVs in patient 340 sample B03. As expected, we find multiple shared SNVs, and two of the three iSNVs in patient 340 sample B03 occur as SNPs in patient 340 sample C03. In Figure 6B, we see seven shared SNPs and four shared iSNVs. All of the iSNVs occur in both patient 639 sample D02 and patient 639 sample G01. These results highlight our ability to identify iSNVs and the feasibility of using iSNVs for identifying paired samples and potential transmission pairs.

**Figure 6:**
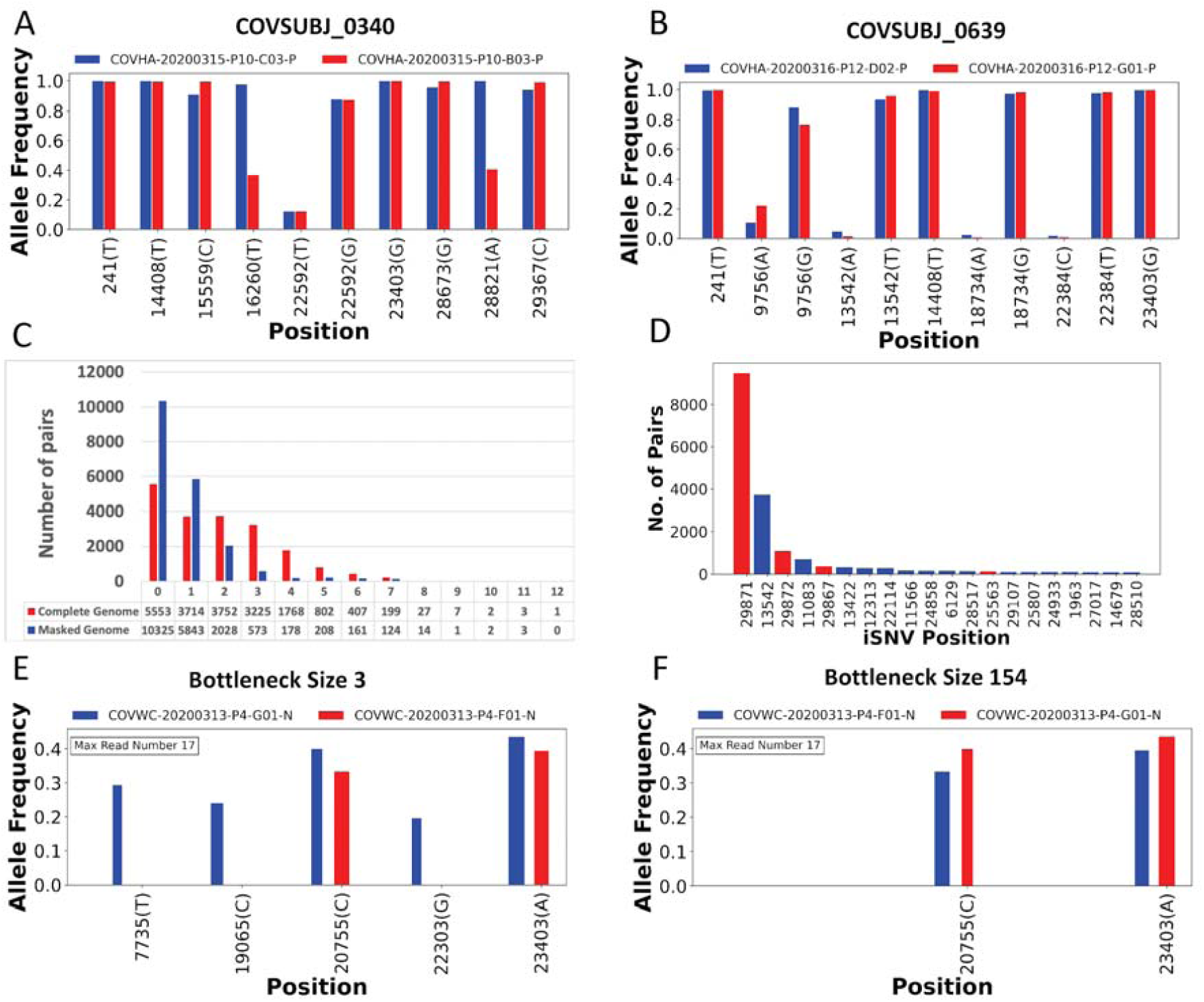
In-depth analysis of shared iSNVs. **A.** Paired samples from patient COVSUBJ 0340 in NYC. **B.** Paired samples from patient COVSUBJ 0639 in NYC. **C.** The distribution of the number of genomic pairs and their shared iSNVs. **D.** The number of samples with iSNVs at given nucleotide positions. Red color represents positions that are highly homplasic and masked in the bottleneck analysis. **E, F.** Allele frequencies and presence of shared iSNVs between two unpaired samples. Blue color represents donor and red color represents recipient. The bar width is proportional to the number of reads supporting the variants. The minimum bar width represents 10 reads. Bottleneck size was estimated to be 3 for **E** and 154 for **F**.

We next calculated the number of shared iSNVs among all possible pairs of NYC samples (Figure 6C). For each pair we consider both possible assignments of donor and recipient, narrowing down the donor alleles to only include those with AF between 0.02 and 0.5, and considering a site to be shared if the recipient also has that same variant present as either a iSNV or SNP. We show these results on the raw data from the iSNV calls, as well as on the same data but after applying masking to sites near the ends of the genome. For the raw data before masking, most pairs have 0 to 3 shared variants, with about 150 pairs having 4 or more shared SNVs (Figure 6c). After masking sites near the genome ends, these numbers drop substantially by reducing likely noise from the variant calls, and we see most pairs sharing 0 to 2 variants. When examining each possible pair, one immediately noticeable trend is that site 29871 yields strong signals for shared SNVs between samples with large and similar AFs. We also observe that the number of samples with a variant at that site is unusually high (Figure 6d). In Figure 6 panels E and F, we see two examples of pairs of samples that not only share multiple iSNVs but also at a similar AF. In these pairs, we find many instances of large estimated bottleneck sizes. The lower estimate of 3 for the pair in Figure 6E is likely due to the variant present at a high AF in the donor at site 7735 that was absent in the recipient.

## Discussion

In this study, we have analyzed over 7,000 SARS-CoV-2 genomes in addition to RNA-seq datasets from 151 COVID-19 positive patients in depth to describe the intrahost variation in SARS-CoV-2. Our analyses yielded four major observations. First, the iSNV mutational spectra closely match the SNP mutational spectra inferred from the consensus genomes. In particular, the SARS-CoV-2 genome is enriched with C>T changes overall, both for iSNVs and SNPs. Genes NSP6 and NSP10 are particularly enriched for T>C mutations, while NSP7 has an enrichment of A>C SNVs. Second, the mutational profile of SARS-CoV-2 largely matches that of other Coronaviruses, but with some key differences. SARS-CoV-2 has a significantly larger proportion of G>T changes in both iSNVs and SNPs, when compared to SARS-CoV-1 and MERS. Additionally, we did not see A>T SNVs in SARS-CoV-1, as previously reported (*31*). Third, while the SV spectra is likely reflecting the transcriptome landscape of SARS-CoV-2, we detected a significant fraction of small indels that fuel the genetic diversity of SARS-CoV-2. Fourth, the mutational spectra of the SNPs and iSNVs indicate that there is a complex interplay between endogenous SARS-CoV-2 mutational processes and host-dependent RNA editing. This observation is in line with several recent studies that propose APOBEC and ADAR deaminase activity as a likely driver of the C>T changes in the SARS-CoV-2 genomes (*14*). Of note, this recent study also reported that the number of observed transversions are compatible with mutation rates found in other Coronaviruses (*8, 14*).

We also reported high sequence conservation within the NSP3 region, a region that is one of the most diverged from SARS-CoV-1 and MERS-CoV. A number of convergent findings suggest de-mono-ADP-ribosylation of STAT1 by the SARS-CoV-2 NSP3 as a putative cause of the cytokine storm observed in the most severe cases of COVID-19 (*32*). The lower mutational complexity of NSP3 agrees with its functional implications in viral replication, and thus the need to conserve its protein structure/function (*31, 33*). Thus, NSP3 may be a good target for drug development since it is well conserved and is essential for viral replication. Follow up studies will be required to solidify functional implications of these observations.

We also investigated the potential impact of iSNVs and SNPs on probes and primers commonly used in RT-PCR based detection and amplicon sequencing of SARS-CoV-2. Most probes we analyzed contain both SNPs and iSNVs. While many platforms can tolerate a few single nucleotide mismatches without the loss of target hybridization, the overall diversity exhibited by SARS-CoV-2 presents potential challenges for probe and primer development. Since we observed a close connection between the SNPs and iSNVs, for future probe and primer designs it could be useful to track the iSNVs to potentially predict and avoid variable regions of the genome. With the integration of these data into design processes at early stages, greater sensitivity could be achieved for hybridization primers and probes even as the virus evolves.

We analyzed paired samples taken from the same COVID-19 positive patient within 24 hours of one another to analyze AFs of SNP and iSNVs. We found that the SNP and iSNV profiles and AFs were concordant, indicating the potential of using shared SNPs and iSNVs and their respective AFs for tracking intrahost SARS-CoV-2 population dynamics. We also scanned all of the NYC COVID-19 positive samples for putative transmission pairs; we highlighted two examples of potential direct or indirect pairs given shared iSNVs at strikingly similar, high AFs. Out of all samples, we found that the majority of pairs show no signal for an inferred large bottleneck. This is to be expected given that the majority of pairs in a large batch of sequenced SARS-CoV-2 samples are not expected to have been direct or indirect transmissions. Of note, the recent report of De Maio *et al. (34)*, many sites were examined that showed extensive homoplasy. While these analyses cannot confirm sample pairs as having been involved in direct transmissions without additional confirmatory metadata, this exploratory analysis suggests the possible presence of such transmission pairs (*29*).

Despite the potential for tremendous insight, the study of intrahost variation in viruses can be confounded by multiple factors. First, the estimated AFs are impacted by variable coverage and transcription patterns. Second, low viral load (Ct values above 32) in samples can have an impact on downstream sequencing and analysis (*35, 36*) (Supplementary Figure 3). Third, previous studies such as De Maio *et al. (34)* highlight SARS-CoV-2 sites marked as prone to high homoplasy and need to be taken into consideration for transmission analyses. Lastly, lack of additional metadata imposes a barrier to an in depth study of transmission events. These factors should be addressed in the future studies of iSNVs in SARS-CoV-2.

In summary, our analysis of intrahost variation across 151 samples from COVID-19 positive patients revealed a complex landscape of within-host diversity that will likely shed additional light on the elusive mechanisms driving the rapid dissemination of SARS-CoV-2. Metatranscriptomic analysis is a powerful tool for interrogating the genomic and transcriptomic landscape of RNA viruses, as it provides a simultaneous peek into viral, bacterial, and host gene expression. Future studies able to integrate all three of these perspectives may hold the key to novel therapies and treatments of this devastating pandemic.

## Materials and methods

### Datasets

We downloaded 6,928 SARS-CoV-2 consensus genomes from the GISAID database, available on April, 18th, 2020. We only selected high quality, complete (>29 Kbp) genomes. We used read data from 11 patient samples collected by Baylor College of Medicine in Houston, Texas. We have also used read data from 140 patient samples collected by Weill Cornell College of Medicine in New York City, New York. Both datasets consist of Illumina NovaSeq 6000 paired-end reads. Host and bacterial genetic material has been removed from the datasets, and we performed all analyses on the viral read data.

For the other coronaviruses data we used 42 samples of SARS-CoV-1 and 53 samples of MERS viral read data (*37*) sequenced by University of Maryland School of Medicine in Baltimore, Maryland.

In total, we analyzed 7,079 SARS-CoV-2, 42 SARS-CoV-1, and 53 MERS samples.

### Read QC and mapping

We processed the Illumina paired-end reads using Trimmomatic ver. 0.39 (*38*) to remove adapter sequences and trim low quality base pairs. We used a universal set of Illumina adapters as a reference for the adapter removal. We set the maximum mismatch count to 2, palindrome clip threshold to 30 and simple clip threshold to 10. We also trimmed leading and trailing low quality (quality value below 3) and ambiguous (N) base pairs. Finally, we applied sliding window trimming cutting the read if the quality score of 4 contiguous bases made the average score drop below 15. After trimming in the final read set we included the reads above the length of 36 with both reads from a pair passing quality control.

We aligned the trimmed reads to the reference genome using Burrows-Wheeler Alignment tool (BWA) ver. 0.7.17 (*39, 40*). We have used paired-end mode for mapping reads to the SARS-CoV-2 reference genome (NC_045512).

We used SAMtools ver. 1.9 to convert the output of *BWA* from SAM to BAM format, and to sort and generate indices for the BAM files (*41*).

### SNV calling and annotation

We used LoFreq ver. 2.1.4 to perform variant calling on the trimmed and mapped reads (*42*). We have filtered the variants with the default LoFreq parameters: minimum coverage was set to 10, phred quality-score set to Q20 (99%), and strand-bias FDR correction p-value is greater than 0.001. We have also filtered out the variants occurring below 0.02 AF threshold for the subsequent analyses, and required all iSNVs to be supported by 10X minimum coverage. We annotated the SNVs found in each of the datasets with snpEff ver. 4.3 (*43*). We used SNPGenie (*44*) with the default set of parameters to estimate the genetic diversity and non-synonymous to synonymous diversity ratios in SARS-CoV-2, SARS-CoV-1 and MERS data.

### SV calling

Structural Variations were identified using Manta (version 1.6.0) (*45*). Subsequently the SV calls were merged using SURVIVOR (v1.0.7) (*46*) using a 100 bp maximum distance between the breakpoints and requiring that the SV types are in agreement in order to merge two SV across the samples. We annotated the SV using a simple 1bp overlap method using bedtools (v2.27.1) (*47*) intersect using the annotations. The same method was used to establish if the start or stop breakpoints of an SV are overlapping with the TRS sites. To test the significance of the overlap we used a permutation test where we randomized the TRS sites (using bedtools random) to generate random TRS with length of 5bp, 1000 times and calculated per TRS the number of start/stop breakpoints of the SV catalog. Subsequently we used this together with the observed overlap using a Kolmogorov–Smirnov (ks.test) with an alternative set to “two.sided” in R (v 3.2.2).

To generate SV and SNV densities we computed the number of variations per type within a 100bp window. For each variant we counted 1/AF where AF is the frequency of that variant across the samples. This was done based on a custom script available on request. The plot was generated using Circos (v 0.69-8) (*48*).

### Phylogenetic tree construction

We used Parsnp (ver. 1.2) (*49*) to align the GISAID genomes. We set the maximal cluster D value to 30,000, and the rest of the parameters were set to the default values. We used RAxML (*50*) to infer a phylogenetic tree from the GISAID alignment. We ran RAxML with default parameters using GTRCAT approximation model for tree scoring. We used the best-scoring maximum likelihood tree output from RAxML.

### Variation in alignment of GISAID assemblies

A multiple sequence alignment of the 6,928 SARS-CoV-2 assemblies was generated with mafft v7.458 (*51*) with the *-auto* option. Variation per column in the alignment was generated with bit v1.8.02 (*52*) which utilizes the scikit-bio (*52, 53*) implementation for calculating Shannon uncertainty.

### Probe and primer mapping

Primer and probe sequences were derived from the WHO website (*54*) and hCoV-2019/nCoV-2019 Version 3 Amplicon Set (*55*). We mapped probes and primers against the SARS-CoV-2 reference genome (NC_045512) with bowtie2 (*56*). Analysis of the primer and probe mapping regions was performed with a custom Python script and visualizations were done with R-3.6.1.

### Transmission Analyses

To compute the number of shared iSNVs in each genomic pair, we utilized the variant calling results. We conducted pairwise genome comparisons and counted the number of shared variants within individual pairs. For each pair, we consider both combinations of one sample as a putative donor and one sample as a putative recipient. Shared iSNVs were then defined as iSNVs that share the same variant nucleotide between the two samples, and where the variant frequencies in the assigned donor sequences are from 0.02 to 0.5. We examined variants with frequencies ≥ 0.02 as the cutoff for conservative estimates to avoid including variants caused by sequencing errors. For the 140 samples from New York, given that we consider each pair twice, there are 19,460 pairs. Note, since we are looking for putative transmission events, we can only consider samples within the same geographic region, so we limited our analyses to the 140 samples that all came from New York. We masked the iSNVs that occur between positions 1-55 and 29804-29903 in the genome. Additionally, we masked 25 nucleotide positions between 56-29804 that are highly homoplasic. These positions are more prone to sequencing and mapping errors (*34*), and therefore were not used in the transmission analyses.

We applied the BB bottleneck software to approximate SARS-CoV-2 bottleneck sizes, that is, the founding viral population size in the recipient host (*57*). Since the variant frequencies in recipient samples partially rely on stochastic replication processes in the early infection, we take all iSNVs (with any AFs) into account (from 0.0 to 1.0) within a shared variant for putative recipients. Furthermore iSNVs from either donors or recipients are supported by at least 10 reads to be included in the bottleneck size analysis. We use the AFs of shared iSNVs between putative donor and recipient pairs as input for the *BB bottleneck* APPROX mode (*57*). If the recipient does not have the iSNV with the same base at the same site as the donor or simply does not have any variant called at position i while mapping to the reference sequence, we assign the recipient a 0.0 AF at that position. Finally, we consider the case where the iSNV base is the same as the reference sequence base. In this case, for instance, when a variant is called at a site with 0.7 AF and no other variants are present, we take the reference base as an iSNV with 0.3 AF if there are no other reads present with an alternate allele and there are at least 10 reads mapping to the reference base.

## Acknowledgments

The authors would like to acknowledge feedback and discussion contributions on the effects of variants on the qRT-PCR detection methods provided by Jamie Purcell. The authors would also like to thank Luay Nakhleh for suggestions specific to comparative genomic analyses of SARS-CoV-1 and MERS-COV. Finally, the authors would also like to thank all members of the COVID-19 International Research Team (www.cov-irt.org) for their helpful feedback during weekly meetings.

## Disclaimer

This material has been reviewed by the Walter Reed Army Institute of Research. There is no objection to its presentation and/or publication. The views and conclusions expressed in this article are those of the authors and do not necessarily reflect the official policy or position of the Department of the Army, Department of the Navy, Department of Defense, ODNI, IARPA, ARO, or US Government.

## Funding information

N.S. and Y.L. are supported by the Department of Computer Science, Rice University. Q.W., D.A., T.J.T, and R.A.L.E. are supported by startup funds from Rice University. M.J. is supported under NIH award No. R01HD091731 from the NICHD. F.J.S. acknowledges funding and part of the data was produced by Baylor College of Medicine under NIAID (U19AI144297-01). A.B. is supported by supplemental funds for COVID-19 research from Translational Research Institute through NASA Cooperative Agreement NNX16AO69A (T-0404) and further funding was provided by KBR, Inc. D.P. is supported by the European Research Council (ERC-617457-PHYLOCANCER), Spanish Ministry of Economy and Competitiveness, and Xunta de Galicia.

## Author contributions

N.S. led the iSNV and SNP analyses, interpreted the results, generated the figures, and wrote the manuscript. M.J. analyzed the impact of polymorphisms on probes and primers, and generated figures.Y.L. analyzed single nucleotide variant data and generated the figures. D.A. analyzed phylogenetic data and generated the figures. M.D.L. analyzed genomic data and generated figures. Q.W. analyzed and interpreted viral transmission data, generated figures, and wrote the manuscript. R.A.L.E. interpreted the viral transmission and phylogenetic data, and wrote and edited the manuscript. S.V. edited the manuscript, provided exchange of ideas, and generated figures. C.M. provided the RNA-seq data and contributed to the manuscript. T.J.T. supervised the analyses, interpreted the data, edited, and wrote the manuscript. M.M. and F.J.S. lead the SV analysis, interpretation of the data and edited and wrote the manuscript. A.B. edited the manuscript and provided exchange of ideas. K.T. reviewed the SNV commands and called variants in public COVID-19 metatranscriptomes for comparison. D.P. proposed some of the analyses, helped with their interpretation, and contributed to manuscript writing. All co-authors read and edited the manuscript and provided constructive feedback.

## Competing interests

Authors declare no competing interests.

## Data availability

Variant calling files, raw data and supplementary figures and files are available at https://rice.box.com/v/SARS-COV-2-SNV-data. Assembled genomes for SARS-CoV-2 used in the analysis are available at GISAID. SARS-CoV-1 and MERS read data were obtained from the study PRJNA233943. Scripts used for data analysis are available at https://gitlab.com/treangenlab/covirt_scripts. Scripts used for probe and primer analysis and visualization are available at: https://github.com/COV-IRT/microbial/tree/master/manuscript_reference

## Notes

### Competing Interest Statement

The authors have declared no competing interest.

