## Supplementary Material for "Hidden genomic diversity of SARS-CoV-2: implications for qRT-PCR diagnostics and transmission"

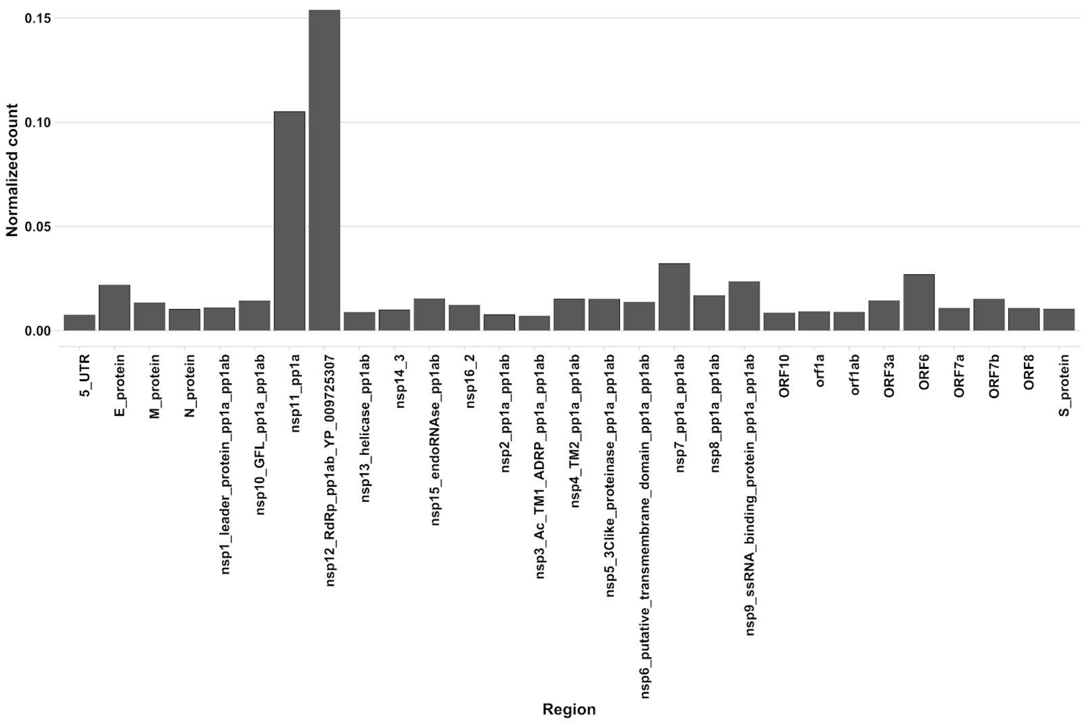


**Supplementary Figure 1:** Structural Variation concentration across annotated regions for SARS-CoV-2. Number of insertion and deletions below 1000bp are counted per region and are normalized by the size of the region showing a clear enrichment in certain parts.

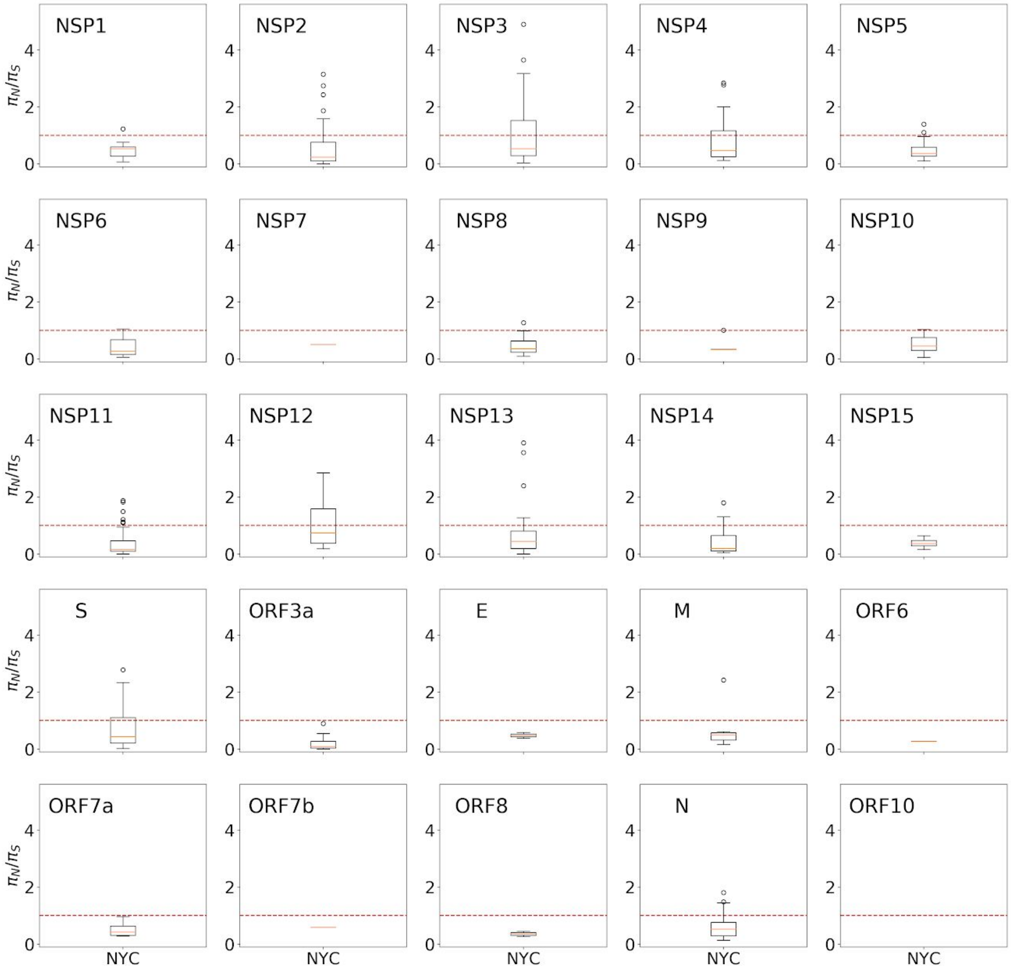


**Supplementary Figure 2:** Non-synonymous to synonymous diversity ratios for iSNVs and SNPs in NYC data mapped onto NSP/ORFs of SARS-CoV-2 genome.

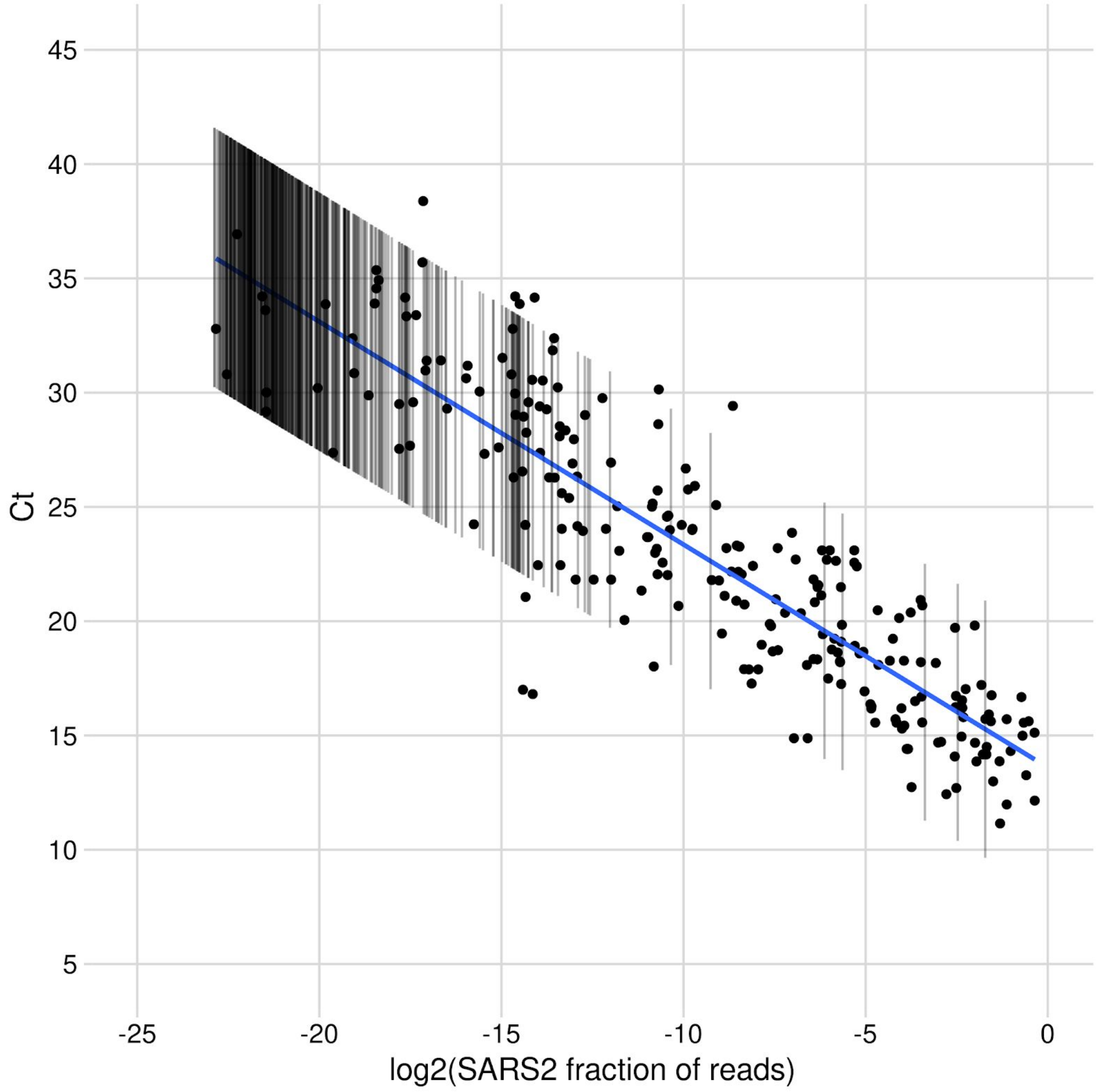


**Supplementary Figure 3:** SARS-CoV-2 mapped read fractions vs Ct. PCR positive samples are shown as points. Lines represent the 95% prediction interval for PCR negative samples with a non-zero number of SARS-CoV-2 mapped reads.

**Supplementary File 1:** WHO detection probes and primers, Artic and Tomer Altman sequencing primers with SNP and iSNV occurrences indicated are summarized in this Excel file.
